# SecretTarget: A pipeline for identifying host-interacting effector candidates through secondary localization features

**DOI:** 10.64898/2026.08.09.743782

**Authors:** Alexander Thomas Julian, Alyssa Britney Barnes, Jean-François Pombert, Jialing Xiang

## Abstract

Intracellular parasites cause over 226 million disability-adjusted life years (DALY) globally each year. Current treatment methods fall short of desirable impact due to adverse effects and growing resistance, shifting focus to parasitic effectors for future therapeutic development. Unfortunately, the divergent nature of parasitic proteins has impeded *in silico* discovery of – and functional inference for – parasitic effectors. Here, we present SecretTarget, a pipeline designed to identify host-interacting effector candidates through secondary localization features. By truncating signal peptides from predicted extracellular proteins (ΔSP), we unmask potential underlying localization features that dictate subcellular trafficking within the host. Applying our pipeline to a set of *Toxoplasma gondii* secreted effectors with known localizations and interactions, we propose a novel host ER-parasite interaction critical for parasite survival, recapitulate published localizations, and provide meaningful biological insights aligning with recent host-parasite interaction discoveries.

## Introduction

Intracellular parasites are a global health burden that most significantly impacts impoverished nations [1]. Conservatively, intracellular pathogens cause ∼226 million disability-adjusted life years (DALY) – a measurement combining years lived with disability and years lost to premature death – globally each year [1]. Across all ages, nearly half (46.0%) of DALY due to intracellular parasites are attributed to children under the age of 5 [1]. Treatment of these pathogens is challenged by growing resistance to staple therapeutics [2,3] and alternatives are plagued by adverse side effects [4,5]. Intracellular parasites are categorized as obligate if a host is necessary for replication (*e.g. Plasmodium falciparum* [6]), or facultative if replication within a host is optional (*e.g. Listeria monocytogenes* [7]). Once internalized, intracellular parasites can reside directly exposed to the host intracellular environment (*Rickettsia prowazekii* [8]), or within a replicative sanctuary, such as parasitophorous vacuoles (PV; *Mycobacterium tuberculosis* [9]) and inclusion bodies (*Chlamydia trachomatis* [10]).

Bound ribosomes are responsible for the translation of the majority of parasitic proteins secreted via the Sec translocon. Though functionally identical to free ribosomes, bound ribosomes are found on the ER membrane and translate proteins destined for secretion, the cell membrane, and cellular trafficking. In contrast, free ribosomes – located freely within the cytoplasm – translate proteins destined to remain in the cytoplasm or targeted to specific compartments by sequence embedded sorting signals, such as the nucleus by the nuclear localization signal (NLS) or the mitochondria by the mitochondria targeting signal (MTS) (Fig. 1A). Proteins destined for secretion are typically labeled – though not strictly required – with a physiochemically conserved signal peptide (SP) sequence at their N-terminus [11,12]. The emergence of the SP from a translating ribosome typically recruits chaperones that bind, halt, and deliver the translation complex to the Sec translocon, initiating translocation across the membrane and resuming protein translation [12]. Eukaryotic Sec-mediated translocation occurs across the ER membrane, requiring intermediate steps for secretion to the extracellular space (Fig. 1A). In contrast, prokaryotic Sec-mediated translocation occurs across the cell membrane, enabling either direct secretion (Gram-positive bacteria) or Type II/V secretion system-mediated secretion (Gram-negative bacteria) to the extracellular space.

**Figure 1.**
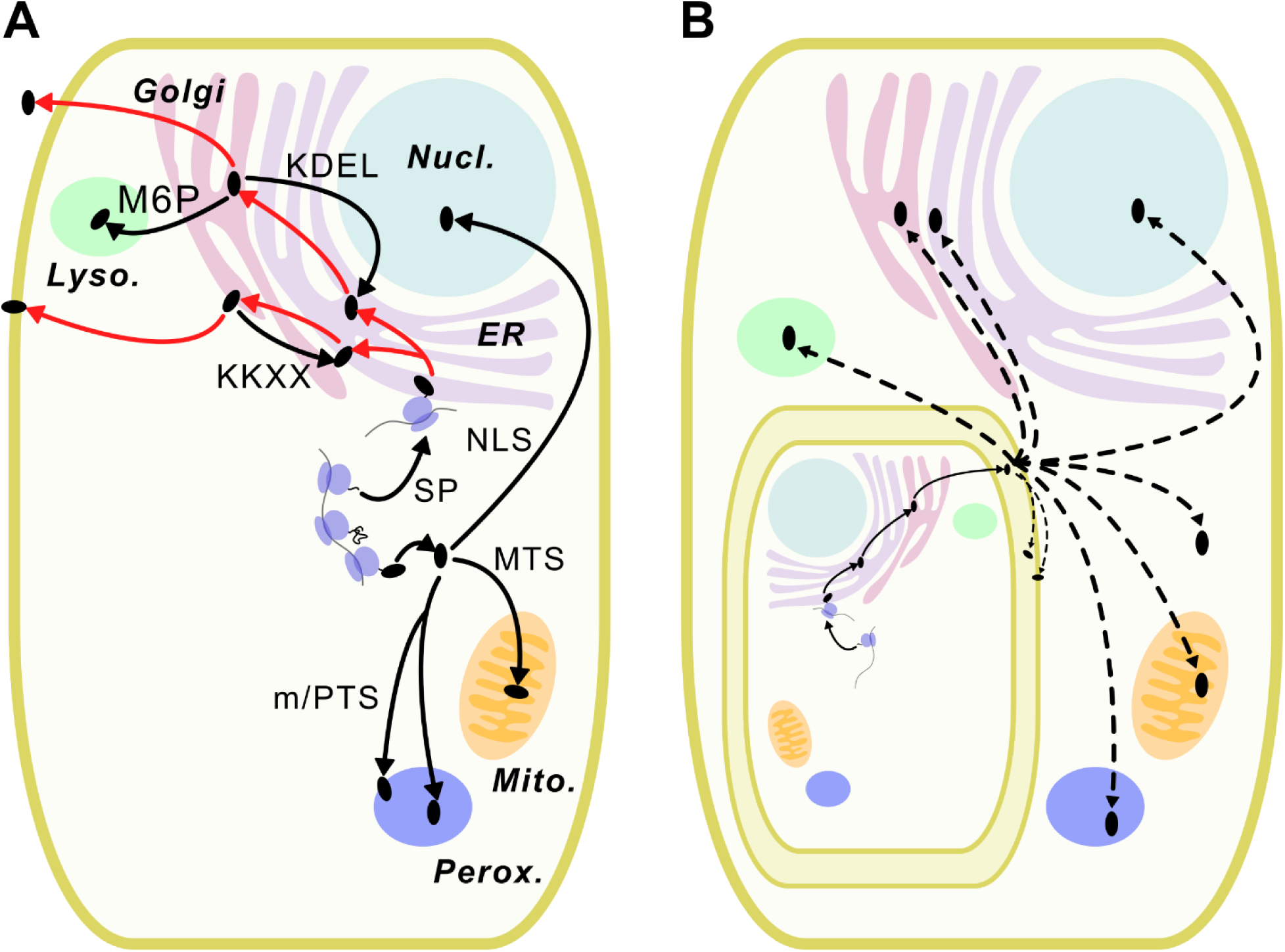
Intracellular parasite protein localization internally and within host. (A) Eukaryotic protein localization is commonly dictated by sequence embedded signals (text next to arrows). Proteins translated by free ribosomes are designated to stay inside the cytosol, unless specific signals direct them to intracellular compartments such as the nucleus (NLS; nuclear localization signal), the mitochondria (MTS; mitochondrial target signal), or the peroxisome (PTS; peroxisomal target signal). Signal peptides (SP) initiate formation of bound ribosomes by recruitment of chaperones that mount the translation complex to the ER, where the nascent protein is translated across the membrane. Once in the ER lumen, proteins are sent to, and processed by, the Golgi Apparatus for secretion to the extracellular space unless additional signals direct them to other compartments, such as M6P to the lysosome, KKXX (Lys-Lys-X-X) to the ER membrane, or KDEL (Lys-Asp-Glu-Leu) to the ER lumen. (B) Encapsulated intracellular parasites secrete their proteins into the parasitophorous space (solid arrows) which can be transported into the host cytoplasm and localize to host organelles (dashed arrows).

For intracellular parasites, the host cytoplasm – or the parasitophorous space (PS) when isolated from the cytoplasm – serves as the “extracellular space” for secreted proteins. Some encapsulated intracellular parasites may use additional mechanisms for transporting secreted proteins across the membrane of their replicative sanctuary and into the host cytoplasm – *e.g.* across the PV membrane (PVM) via the PTEX system in *P. falciparum* [13]. Secreted proteins that establish a replicative niche by modulating host behavior, hijacking host machinery, and requisitioning host nutrients and metabolites are collectively termed effectors. Disruption of effector functionality diminishes virulence, parasite growth, and proliferation, rendering them prime candidates for novel therapeutic interventions [14–17]. However, the host-pathogen evolutionary arms race has driven significant divergence in parasitic protein sequences, obstructing *in silico* efforts to elucidate function by sequence homology [18]. Though numerous effector-related tools and pipelines have been developed, they tend to be narrow in scope [19], focus solely on identification without addressing effector interaction or function [19–21], and in some cases, are no longer accessible [22].

Proper functionality and interaction with the host requires accurate cellular trafficking of effectors to specific host compartments (Fig. 1B) – likely facilitated by host transport machinery to overcome the spatial constraints of diffusion-based localization. This co-opting of existing host pathways implies the existence of “secondary” localization (SL) features (explicit and implicit, *e.g.* MTS and structural folds, respectively) within effectors that are recognized by host trafficking networks. Consequently, deciphering the host-side subcellular localization of secreted parasitic proteins within the host can help demystify their molecular mechanisms and highlight promising effector candidates for therapeutic targeting. For nearly half a century, significant effort has been dedicated to identifying signal peptides and predicting protein localization directly from protein sequences [23], with modern state-of-the-art predictors utilizing deep learning [24–26] making remarkable strides beyond foundational methods [27]. To bridge the gap between general protein sorting and host-parasite interactions governed by these secondary signals, here we present SecretTarget, a pipeline for identifying host-interacting effector candidates through secondary localization features.

### Pipeline Theory and Overview

The SP is the first feature translated by the ribosome and recognized by chaperones, thus ER localization and secretion – in eukaryotes and prokaryotes, respectively – takes priority over any other features potentially present within the protein. Specific features work synergistically with SPs for proper localization (Fig 1A), while targeting potential of other features would be completely abolished. Because many localization features require cytosolic chaperones (*e.g.* MTS, NLS, PTS), concealment within the ER prevents proper localization, and thus is atypical to be present alongside SPs. While detrimental to standard cellular trafficking, the failure of SL feature recognition within the ER can be extorted by intracellular parasites to secrete effectors with host-targeting localization signals.

Simply, localization predictors utilize a two-step system: detection of localization features and localization classification. Each localization feature has a detector responsible for indicating the likelihood of being present in the provided protein. The classifier receives all feature likelihood signals, assigning each an amplification factor (referred to as a weight). The classifier weighs all incoming signals and predicts the most likely localization. Because SPs are the first localization feature that chaperone proteins encounter, the classifier assigns significant weight to SPs when identified, masking other features (Fig. 2A). In the case of intracellular effectors, the SP signal will computationally bury SL feaures. To overcome this difficulty, we developed SecretTarget, a pipeline designed to unmask previously hidden SL features by truncating SPs from predicted secreted proteins (Fig. 2B and C).

**Figure 2.**
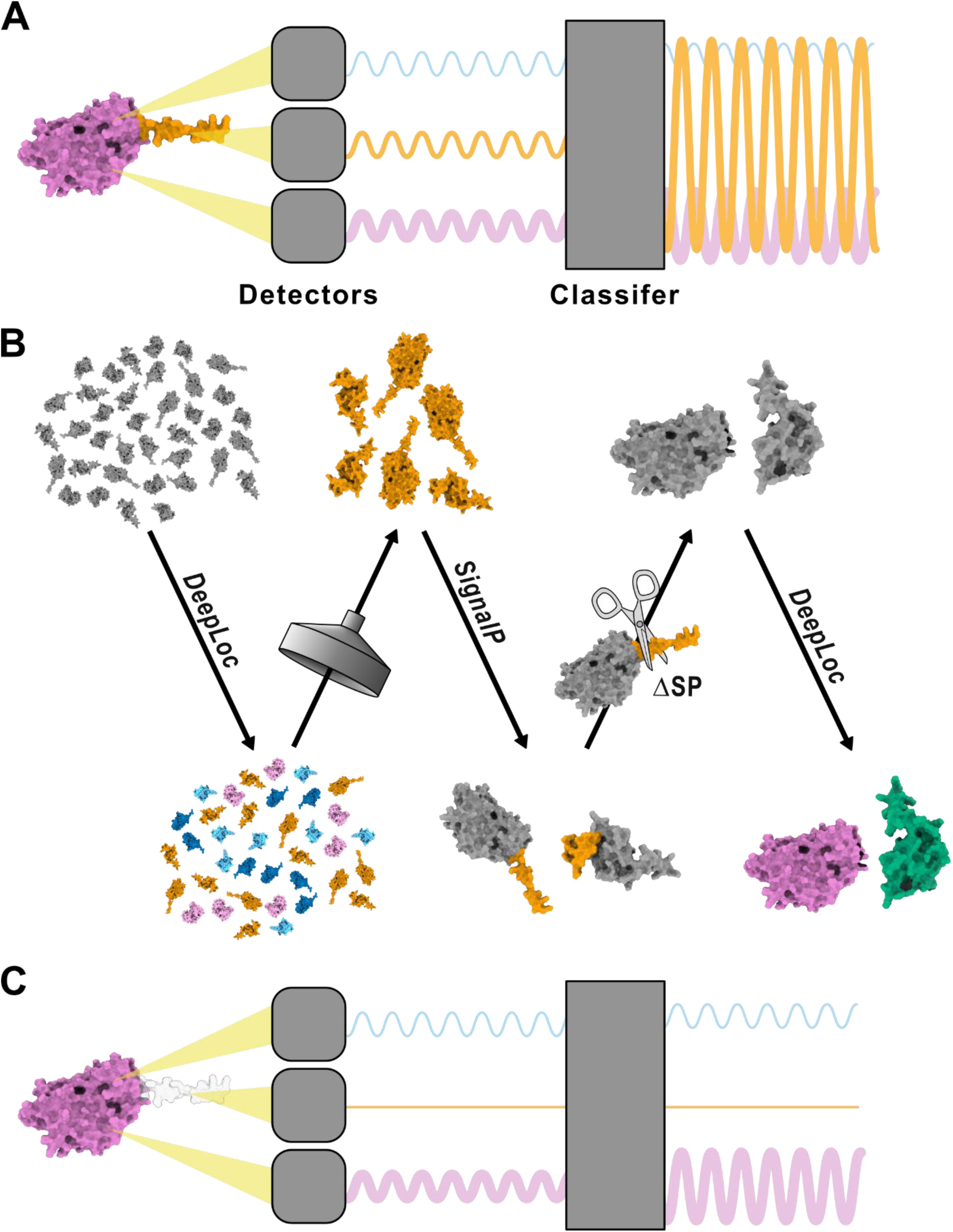
SecretTarget theory and pipeline overview. (A) Simplified representation of a localization predictor. Detectors check the protein for the specific features and send signals to the classifier communicating the likelihood of the feature being present. The classifier applies an amplification factor to each feature and predicts a localization after considering all adjusted signals. In cases where proteins contain underlying signals (purple) in addition to an SP (orange), the significance factor applied to the SP signal overwhelms, and masks, all other signals. (B) Initial protein localizations are predicted using DeepLoc and non-extracellular proteins are filtered from the dataset. Protein signal peptides (SP) are identified with SignalP and are truncated from protein sequences (ΔSP); proteins without an identified SP are removed from the dataset. Finally, putative host subcellular localizations for the remaining proteins are predicted again using DeepLoc. (C) Similar to (A), however, SecretTarget has been run on the input protein, removing its SP (ΔSP) and unmasking the underlying localization signal.

First, all proteins are subjected to subcellular localization prediction by DeepLoc (DeepLoc2 and DeepLocPro for eukaryotes and prokaryotes, respectively); proteins not categorized as extracellular are discarded. Next, proteins are interrogated for SPs by SignalP; extracellular proteins lacking an identifiable SP are considered false positives and discarded. Following SP recognition, each protein is subjected to *in silico* truncation to remove the predicted SP (ΔSP), mimicking biological cleavage. Finally, an additional round of subcellular localization prediction is performed on the ΔSP proteins using DeepLoc to reveal their putative SL. SecretTarget operates on a simple multifasta input, removes SPs from secreted proteins, identifies potential underlying SL features, and provides results in an easy to interpret graphic that can be adjusted from the command-line.

### Software and Hardware Requirements

SecretTarget is freely available under the AGPL license and can be downloaded from GitHub (https://github.com/ATJulian-Lab/SecretTarget) or installed using the python pip package manager. SecretTarget was developed and tested on Fedora Linux (v.42/43). The pipeline is dependent on DeepLoc (v.2.1) and SignalP (v.4.1g and/or v.6.0i), which require licenses to download (detailed instructions for obtaining a license, installation, and environment setup are available on GitHub). Because SecretTarget is built around deep-learning software, a high-speed multi-cored computer processing unit (CPU) is recommended alongside a Compute Unified Device Architecture (CUDA)-enabled graphics processing unit (GPU), though not required.

### Case Study – Toxoplasma gondii

*Toxoplasma gondii* is a human-infecting intracellular parasite with a documented seroprevalence up to 90% in some human populations [28]. Congenital toxoplasmosis, the vertical transmission of *T. gondii* from mother to fetus, is considered a public health hazard as fetal infection can result in significant developmental defects and stillbirth [29,30]. While previously only a major concern for the immunodeficient, recent documentation indicates a novel risk to the immunocompetent, notably possible long-term neuropsychiatric and cognitive disorders (such schizophrenia and Alzheimer’s, respectively) [31–34]. Presently no therapeutics exist for latent *T. gondii* infection, with treatments for prophylaxis and acute infection requiring long dosing periods, causing significant adverse events, and exhibiting indicators of decreasing efficacy [35–38].

To analyze the *T. gondii* proteome (Strain: GT1, VEuPathDB Version: ToxoDB-68 [39]) for candidate effector proteins, we ran SecretTarget, specifying SignalP4.1 for its lower stringency to mitigate the loss of SP discovery due to sequence divergence. Of 8640 proteins, initial localization prediction classified 760 as extracellular (Fig. 3A and B). SecretTarget identified and truncated SPs from 484 putative secreted effectors (ΔSP), 323 of which revealed SLs (Fig. 3A and C). The majority of predicted SLs were to the cytoplasm (198), the nucleus (66) and the cell membrane (64), with the remaining split amongst the mitochondria (12), the ER (10), plastid (7), the Golgi (3), and lysosome/vacuole (1) (Fig. 3C). A small portion of proteins maintained extracellular classification while also being assigned an additional SL (24/484), while one third were not reclassified from the extracellular compartment (161/484) despite SP removal (Fig. 3C).

**Figure 3.**
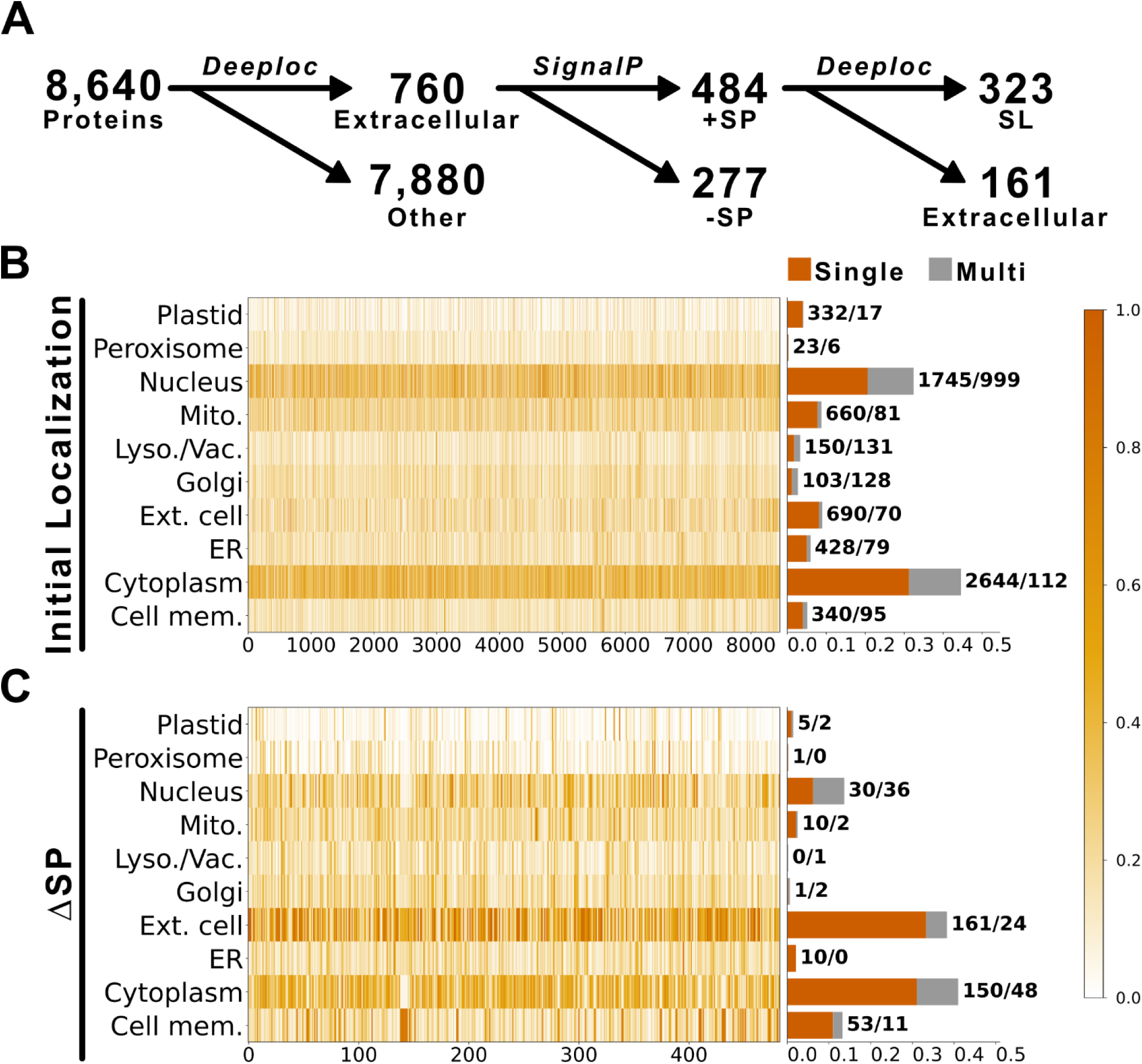
Initial and secondary predicted localizations for the *T. gondii* GT1 proteome. (A) Number of proteins present after each step in SecretTarget. (B) DeepLoc predictions for the 8640 proteins in the proteome; scores are displayed as a heatmap (x-axis: protein position in the dataset). Number of proteins assigned to each compartment is shown as a bar chart; orange indicates proteins assigned to a single compartment, and grey indicates proteins assigned to multiple compartments. Proteins per bar segment are shown to the right in bold. (C) DeepLoc secondary localization predictions for proteins initially predicted as extracellular after *in silico* truncation to remove SPs. *Colorbar indicates scores in heatmaps*.

A curated list of *T. gondii* effectors with known host localizations were [40] used to validate the SL prediction accuracy of SecretTarget. Due to the unique nature of *T. gondii* rhoptry (ROP) and dense granule (GRA) protein delivery, 11 of the 22 sequences were not predicted to localize to the extracellular compartment (Fig. 4A and B). Of the proteins predicted to localize to the extracellular compartment, the majority contained an identifiable SP (Fig. 4A and C; Table 1), enabling SL prediction. Only ∼27% (3/11) of SL predictions match the experimentally determined localizations exactly. However, when factoring in membrane association type and pathogen-host interaction dynamics, all predicted SLs carry biologically relevant insights consistent with SecretTarget results (Table 1).

**Figure 4.**
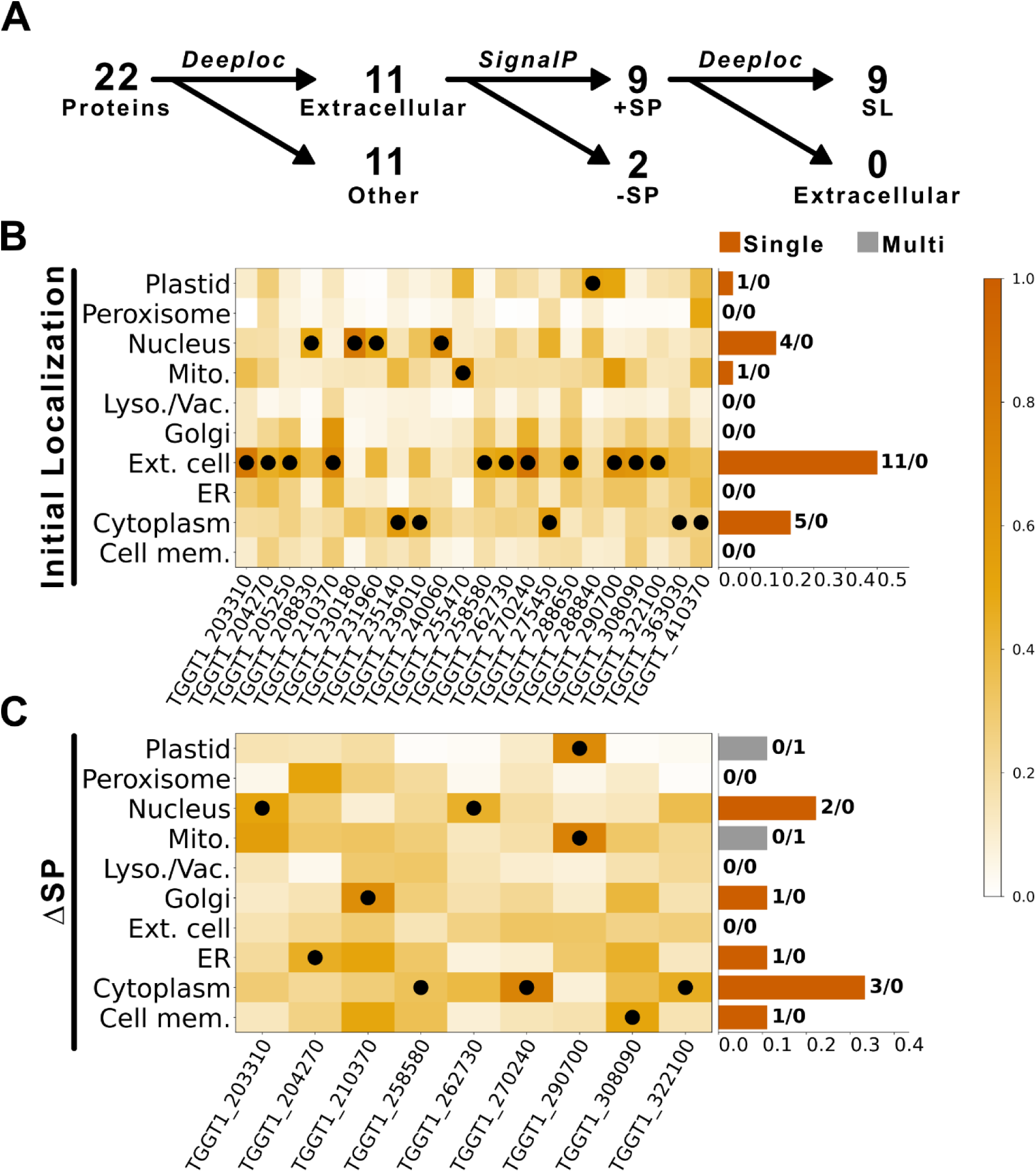
Initial and secondary localizations of *T. gondii* effectors with known host localizations. (A) Number of proteins present after each step in SecretTarget. (B) DeepLoc predictions for 22 proteins with known host localizations; scores are displayed as a heatmap (x-axis: protein position in the dataset). Number of proteins assigned to each compartment is shown as a bar chart; orange indicates proteins assigned to a single compartment, and grey indicates proteins assigned to multiple compartments. Proteins per bar segment are shown to the right in bold. (C) DeepLoc secondary localization predictions for proteins initially predicted as extracellular after *in silico* cleavage of SP. *Colorbar indicates scores in heatmaps*.

**Table 1.** SecretTarget results for curated *T. gondii* effectors.

| Accession # | Annotation | SP | Predicted SL | Published SL | Result<sup>1</sup> | Reference |
| --- | --- | --- | --- | --- | --- | --- |
| TGGT1_203310 | GRA7 | + | Nucleus | PVM | C | [40] |
| TGGT1_204270 | GRA60 | + | ER | PVM | C | [39] |
| TGGT1_205250 | ROP18 | - | - | PVM/ER | - | - |
| TGGT1_210370 | ROP54 | + | Golgi | PVM | C | [41] |
| TGGT1_258580 | ROP17 | + | Cytoplasm | PVM | B | - |
| TGGT1_262730 | ROP16 | + | Nucleus | Nucleus | A | - |
| TGGT1_270240 | MAG1 | + | Cytoplasm | PVM/Cyto. | A | - |
| TGGT1_288650 | GRA12 | - | - | PVM/IVN | - | - |
| TGGT1_290700 | GRA25 | + | Mito/Plastid | PV | C | [42] |
| TGGT1_308090 | ROP5 | + | Cell mem. | PVM | A | - |
| TGGT1_322100 | ROP38 | + | Cytoplasm | PVM/IVN | B | - |

*SL-Secondary Localization; PVM-Parasitophorous Vacuole Membrane; IVN Intravacuolar Network* *A: Predicted localization matches published host destination; B: Predicted localization represents the adjacent soluble compartment exposed to the host (e.g., Cytoplasm for PVM-associated peripheral effectors); C: Predicted localization reflects a documented host organelle contact site, interaction partner, or secondary recruitment site.*

It is well established that the replicative sanctuaries of many intracellular parasites – including *T. gondii* – intimately associates with host organelles, driven by effector anchoring (*e.g.* MAF1 [41]). In *T. gondii*, the organellar association with the host ER is used to modulate host immunity and apoptosis by activating the unfolded protein response [41], despite the host ER membrane acting as a reservoir for Immunity-Related GTPase family protein 6 (Irga6), a host defense that can neutralize *T. gondii* proliferation within hours of infection by disrupting the PVM [42]. Thus, the intimate association of the *T. gondii* PVM with the host ER seems completely counterintuitive, begging the question, how does *T. gondii* avoid destruction while contacting the host ER?

Using SecretTarget, we identified an effector – GRA60 – with a predicted SL of the ER (Table 1). GRA60 – a single pass transmembrane protein found on the cytosolic side of the PVM – is responsible for neutralizing Irga6 [43]. This predicted ER localization suggests that GRA60’s presence at the host ER-PVM interface is a key mechanism for parasite survival, highlighting it as an exciting candidate for novel therapeutic development.

## Conclusion

Intracellular parasites are a significant global health burden, capable of causing illness, chronic disability, and premature death. Because parasitic effectors modulate host cellular behavior to establish a protected proliferative niche, their disruption and inhibition cause substantial degradation in parasitic amplification, making them sought after drug candidates. Unfortunately, significant sequence divergence obfuscates standard *in silico* functional elucidation of parasitic proteins, necessitating the development of novel computational approaches.

Here, we describe SecretTarget, a pipeline for identifying host-interacting effector candidates through secondary localization features. Applying our method to a curated dataset of *T. gondii* ROP and GRA proteins, we recapitulate known interactions in addition to revealing a likely mechanism for the survival of *T. gondii* during close association with the host. The provided case study highlights the potential of SecretTarget to assist in identifying candidate parasitic effectors, providing a unique avenue for uncovering novel drug candidates.

## Author Contributions

A.T.J. conceived the project; A.T.J.; J-F.P. conceptualized the workflow; A.B.B. prototyped the workflow; A.T.J. developed the pipeline; A.T.J. wrote the manuscript; A.T.J.; A.B.B; J.X. reviewed and edited the manuscript.

## List of Abbreviations

DALY: Disability-Adjusted Life Years
ER: Endoplasmic Reticulum
GRA: Dense Granule
MTS: Mitochondria Targeting Signal
NLS: Nucleus Localization Signal
PS: Parasitophorous Space
(m)PTS: (membrane) Peroxisome Targeting Signal
PV(M): Parasitophorous Vacuole (Membrane)
ROP: Rhoptry
SL: Secondary Localization
SP: Signal Peptide

